# RNAi-mediated rheostat for dynamic control of AAV-delivered transgenes

**DOI:** 10.1101/2022.09.30.510386

**Authors:** Megha Subramanian, James McIninch, Ivan Zlatev, Mark K. Schlegel, Charalambos Kaittanis, Tuyen Nguyen, Saket Agarwal, Timothy Racie, Martha Arbaiza Alvarado, Kelly Wassarman, Thomas S. Collins, Tyler Chickering, Christopher R. Brown, Karyn Schmidt, Adam B. Castoreno, Svetlana Shulga-Morskaya, Elena Stamenova, Kira Buckowing, Daniel Berman, Joseph D. Barry, Anna Bisbe, Martin A. Maier, Kevin Fitzgerald, Vasant Jadhav

## Abstract

Adeno-associated virus (AAV)-based gene therapy could be facilitated by the development of molecular switches to control the magnitude and timing of expression of therapeutic transgenes. RNA interference (RNAi)-based approaches hold unique potential as a clinically proven modality to pharmacologically regulate AAV gene dosage in a sequence-specific manner. We present a generalizable RNAi-based rheostat wherein AAV transgene expression is silenced using the clinically validated modality of chemically modified short interfering RNA (siRNA) conjugates or vectorized co-expression of short hairpin RNA (shRNA). For transgene induction, we employ REVERSIR technology, a synthetic high-affinity oligonucleotide complementary to the siRNA or shRNA guide strand to reverse RNAi activity and rapidly recover transgene expression. For potential clinical development, we report potent and specific siRNA sequences that may allow selective regulation of transgenes while minimizing unintended off-target effects. Our results establish a conceptual framework for RNAi-based regulatory switches with potential for infrequent dosing in clinical settings to dynamically modulate expression of virally-delivered gene therapies.

## INTRODUCTION

Adeno-associated virus (AAV) vectors have emerged as the leading platform for most *in vivo* gene therapy applications with potential as a years-if not life-long disease treatment option following a single dose.^1^ However, one key challenge that has become evident from recent clinical trials of systemic AAV-mediated gene therapy is the wide interindividual variability in therapeutic protein expression at the same vector dose, which could lead to phenotoxicity at supraphysiological transgene levels in some cases.^2, 3^ This, along with the difficulty extrapolating the therapeutically efficacious dose range in humans from preclinical data, underscore the need for clinically translatable approaches to modulate transgene expression after AAV administration.^4–6^

RNA interference (RNAi) is an evolutionarily conserved mechanism in which endogenous [microRNA (miRNA)] or exogenous [short interfering RNA (siRNA), short hairpin RNA (shRNA)] short non-coding RNAs downregulate gene expression of mRNA transcripts in a sequence-dependent manner.^7^ As a native pathway that leverages an efficient cellular catalytic mechanism, RNAi can achieve robust, durable, and specific silencing of gene transcripts of interest. In recent years, several RNAi-based drugs have been successfully validated in clinical studies, with demonstrated benefit at low and infrequent doses. Novel delivery solutions along with highly chemically modified RNAs have improved potency, durability, and safety and thus greatly expanded the reach of RNAi therapeutics, culminating in five approved drugs and several others in clinical development.^9–11^ In liver, infrequent delivery of metabolically stabilized siRNA conjugated to N-acetylgalactosamine (GalNAc) results in potent silencing of gene expression that persists for months in humans with favorable safety and tolerability profiles.^12–14^ Recent work has also broadened the scope for siRNA delivery to extrahepatic tissues, with conjugation of 2’-*O*-palmityl (C16) demonstrating wide distribution and durable target knockdown across cell types in central nervous system, eye, and lung.^15^ All of these advances for therapeutic silencing of endogenous disease-associated genes via RNAi also hold the potential for on-demand regulation of exogenously-delivered transgenes in a therapeutic setting.

Given the small footprint of RNAi sequences, complementary binding sites (typically 19-23 nucleotides) for an siRNA or shRNA may be readily incorporated into viral genomes, typically within the 3’ UTR of the vector-encoded transgene. AAV incorporation of binding sites for tissue-specific microRNAs has been exploited to improve specificity of transgene targeting by selectively attenuating expression in undesired cell types.^16–18^ Prior designs for RNAi-based on-switches have leveraged ligand binding to control the processing of shRNAs delivered alongside the therapeutic transgene or to modulate accessibility of endogenous microRNAs to their cognate binding sites on virally-delivered mRNAs.^19–24^ However, their applicability has been limited due either to low dynamic range for modulation, off-target risks, or a lack of validation in the clinical setting.^19^ In contrast, supplying chemically stabilized siRNAs exogenously overcomes the reliance on endogenous miRNAs and offers precise and flexible control of dosage. As an alternative to siRNAs that require repeated, though infrequent administration, RNAi via shRNAs that can be stably introduced into AAV vectors in a gene therapy setting allow continuous regulation of the expressed transgene in *cis* as a single treatment. For AAV-delivered RNAi, embedding artificial shRNA stem sequences within the native scaffold of endogenous microRNA transcripts downstream of weaker Pol-II promoters has been demonstrated to trigger potent knockdown, while mitigating risks related to toxicity, off-targeting, and genomic truncation during viral replication.^25–28^ Prior studies have shown that shRNAs can be expressed at high levels alongside therapeutic transgenes in a single AAV vector in the context of long-term gene complementation for treatment of toxic gain-of-function diseases.^24, 29^

While exogenous RNAi modalities may enable low basal expression of transgenes, the versatility of these systems would be improved if transgene silencing can be reversed to control expression in the on-state. We recently reported a highly potent and generalizable approach for *in vivo* control of RNAi pharmacology using a short, synthetic single-stranded oligonucleotide known as REVERSIR.^30^ REVERSIR molecules functionally abrogate RNAi activity by acting as synthetic high-affinity decoys to sequester RNA-induced silencing complexes (RISC) loaded with complementary siRNA antisense (guide) strands in competition with siRNA target mRNAs. REVERSIR binding thereby prevents RISC-mediated recognition and degradation of target mRNA transcripts and consequently allows their translation. The development of REVERSIR as an antidote for RNAi activity represents a valuable tool that may be co-opted to regulate on-states of exogenously delivered transcripts by enabling induction of transgene expression from RNAi-regulated AAV vectors. Blockade of functional RISC complexes with synthetic complementary oligonucleotides has also been used to inhibit the activity of specific disease-associated endogenous microRNAs, with early clinical data showing translation in humans.^31–35^

Here, we investigate the potential utility of combining RNAi-mediated knockdown with REVERSIR-enabled rescue of transgene expression as a rheostat for AAV-delivered genes. Using hepatotropic recombinant AAV8 (rAAV8) vectors as a model, we assessed exogenous siRNA delivery and viral co-expression of shRNA alongside the transgene as approaches for silencing of transgene expression and demonstrate efficient upregulation of transgene levels by blocking RNAi activity of siRNA/shRNA with REVERSIR technology. Whereas the standard high stability REVERSIR design enabled long-term transgene induction, a less metabolically stable chemical template enabled a shorter duration of transgene upregulation and consequent resumption of RNAi-mediated silencing. Finally, we describe novel and highly specific siRNA sequences that may be employed for clinical development to modulate expression from exogenous vector-delivery systems without causing undesired off-target gene silencing within the endogenous transcriptome of humans and mammalian preclinical models. Together, our studies support the development of RNAi and related REVERSIR technology for regulation of AAV transgenes, particularly for temporal control and dosage refinement in the event of highly variable transduction efficiency in the clinical setting.

## RESULTS

### REVERSIR-mediated induction of transgene under control of vectorized shRNA

We first evaluated potential of a single agent approach for rAAV transgene control whereby silencing of expression may be achieved by co-expression of an shRNA *in cis* from the rAAV vector, with induction then conferred by exogenous delivery of REVERSIR **(Figure 1A).** To explore this possibility, we initially examined the ability of REVERSIR to rescue RNAi-mediated silencing triggered by two well-characterized miRNA-embedded shRNA scaffolds using luciferase assays in cultured cells. We assessed *in trans* repression by co-transfecting transthyretin (*Ttr*)-targeting shRNA or control non-targeting (NT) shRNA embedded within an optimized human miR-30 backbone termed miR-30E or a mouse miR-33 **(Supplementary Table 1)** primary microRNA (pri-miRNA) scaffold along with a luciferase vector containing a fully complementary murine *Ttr* target element within the 3’ UTR.^25, 27^ Expression of TTR shRNA from both scaffolds resulted in significant suppression of luciferase reporter activity relative to the NT shRNA. In both cases, dose-dependent recovery of luciferase reporter expression was observed with co-transfection of 22-mer TTR-REVERSIR but not with a control non-targeting REVERSIR (NT REVERSIR) **(Table 1)** of the same length and chemistry **(Supplementary Figure 1A and Supplementary Figure 1B).**

**FIGURE 1:**
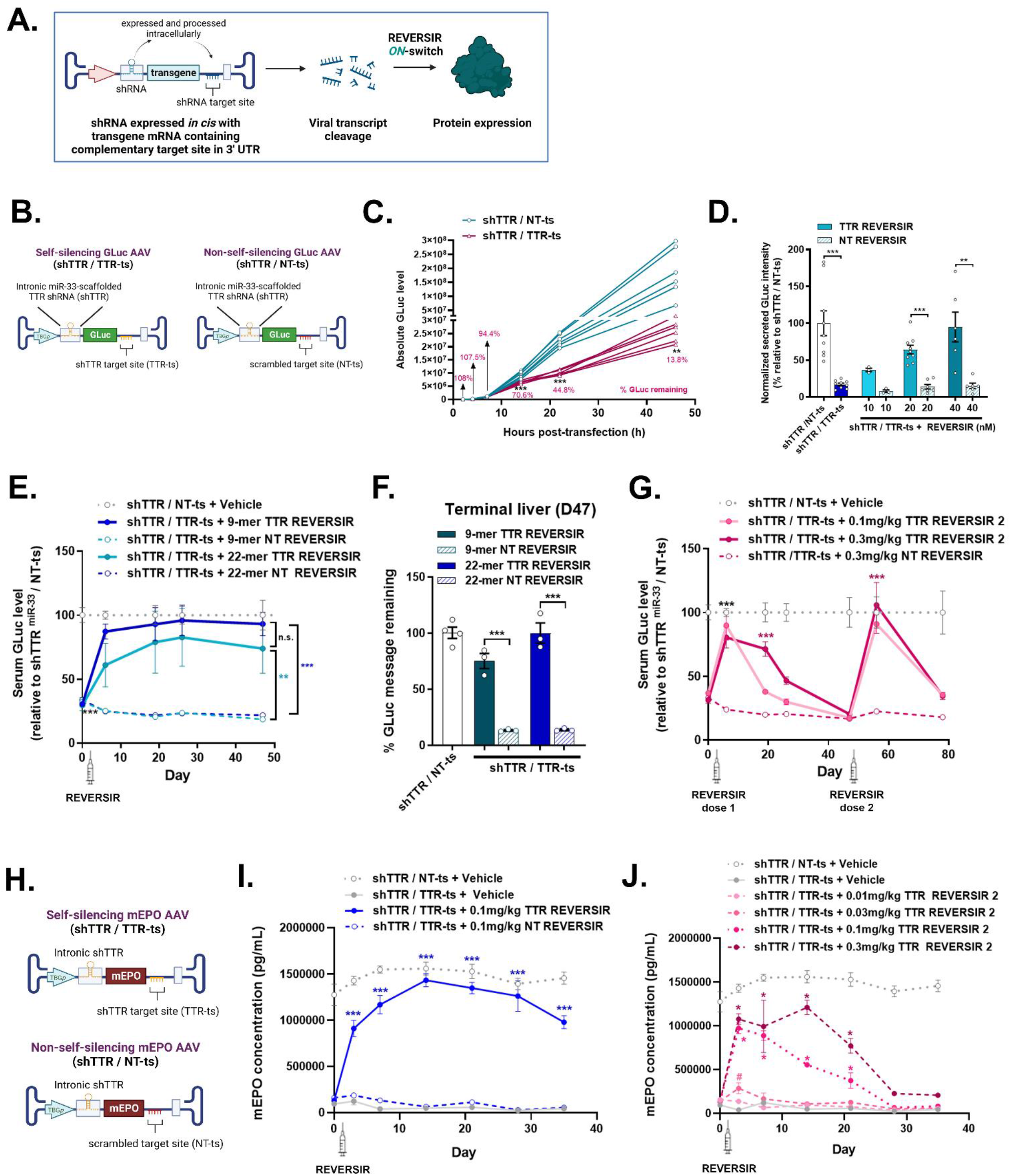
Transgene induction from shRNA-regulated self-silencing rAAV vector using REVERSIR. **(A)** Diagram illustrating self-silencing rAAV vector using intronically-encoded shRNA for transgene silencing and exogenously-administered REVERSIR for transgene induction. **(B)** Viral genome schematic of rAAV8 GLuc reporter vector with intronic TTR shRNA and complementary (‘self-silencing’ TTR shRNA/TTR-ts) or scrambled 3’ UTR target site (‘non-self-silencing’ shTTR/NT-ts) **(C)** HepG2 cells were transfected with the indicated AAV plasmids and media was collected at each timepoint for quantification of secreted GLuc levels. Media was fully exchanged at every collection, with each line corresponding to accumulated GLuc from a single well since prior timepoint. **(D)** Validation of transgene self-suppression and induction with REVERSIR in HepG2 cells. rAAV constructs described in (B) encoding GLuc transgene and intronic miR-33 shRNA along with a Firefly luciferase (FLuc) internal control plasmid were co-transfected with indicated concentrations of 22-mer TTR REVERSIR or NT REVERSIR control. 48h post-transfection, GLuc and FLuc intensities were assayed in cell culture supernatant and lysate, respectively, and GLuc/FLuc ratios computed to normalize for transfection efficiency. **(E - G)** Mice were injected with 2 × 10^11^ genome copies (GC) of rAAV vectors in **(B)** 2 weeks prior to molar equivalent dosing of 9-mer (0.1 mg/kg) or 22-mer (0.2mg/kg) TTR or NT REVERSIR on Day 0 (D0). **(E)** Secreted GLuc levels measured in serum collected prior to and 6, 19, 26, and 47 days following REVERSIR treatment. **(F**) qRT-PCR analysis showing GLuc transcript levels in liver tissue at terminal day 47 timepoint, compared to endogenous *Gapdh* control and plotted relative to shTTR/NT-ts condition set to 100%. **(G)** Longitudinal quantification of serum GLuc levels in mice administered with 0.1mg/kg or 0.3mg/kg 9-mer TTR REVERSIR 2 or NT REVERSIR on D0, followed by a second dose on D47. Bleeds were performed at the same timepoints as in (E), along with additional sample collections on D56 and D78. **(H and J)** Mice were injected with 2 × 10^11^ GC of AAV8 virus encoding mouse EPO (mEPO) transgene under control of TTR shRNA with intact (TTR-ts) or scrambled binding site (NT-ts) in 3’ UTR. **(H)** Vector genome schematics for rAAV encoding self-silencing and non-self-silencing mEPO transgene. Serum EPO concentrations were measured via ELISA on D3, D7, D14, D21, D28, and D35 **(I)** post-treatment with 0.1 mg/kg 9-mer TTR or NT REVERSIR on D0, or **(J)** increasing doses of TTR REVERSIR 2 (0.01, 0.03, 0.1, or 0.3mg/kg) compared to vehicle control on D0. # indicates adjusted *p*<0.05 compared to shTTR / TTR-ts + Vehicle condition by multiple unpaired *t*-test (one per row) corrected for multiple comparisons by Holm-Sidak method. All error bars represent s.e.m.

**Table 1:**
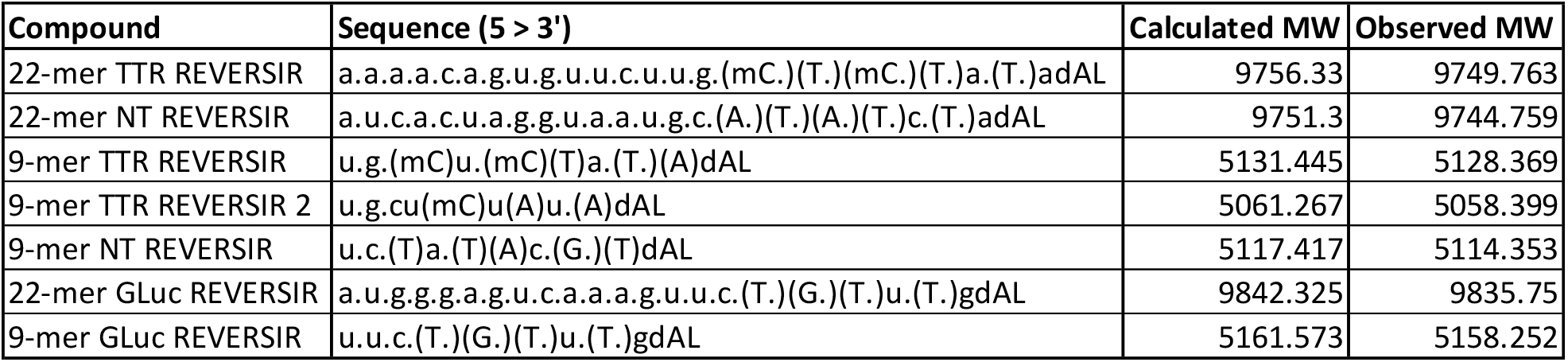
REVERSIR molecules used this study. Legend for chemical modifications is as follows: lower case nucleotides – 2’-*O*-methyl (OMe); upper case nucleotides in parentheses – locked nucleic acid (LNA); (mC) – 5-methyl-cytidine-LNA; dA – 2’deoxyadenosine; L – tri-*N*-acetylgalactosamine.

| Compound | Sequence (5' > 3') | Calculated MW | Observed MW |
| --- | --- | --- | --- |
| 22-mer TTR REVERSIR | a.a.a.a.c.a.g.u.g.u.u.c.u.u.g.(mC.)(T.)(mC.)(T.)a.(T.)adAL | 9756.33 | 9749.763 |
| 22-mer NT REVERSIR | a.u.c.a.c.u.a.g.g.u.a.a.u.g.c.(A.)(T.)(A.)(T.)c.(T.)adAL | 9751.3 | 9744.759 |
| 9-mer TTR REVERSIR | u.g.(mC)u.(mC)(T)a.(T.)(A)dAL | 5131.445 | 5128.369 |
| 9-mer TTR REVERSIR 2 | u.g.cu(mC)u(A)u.(A)dAL | 5061.267 | 5058.399 |
| 9-mer NT REVERSIR | u.c.(T)a.(T)(A)c.(G.)(T)dAL | 5117.417 | 5114.353 |
| 22-mer GLuc REVERSIR | a.u.g.g.g.a.g.u.c.a.a.a.g.u.u.c.(T.)(G.)(T.)u.(T.)gdAL | 9842.325 | 9835.75 |
| 9-mer GLuc REVERSIR | u.u.c.(T.)(G.)(T.)u.(T.)gdAL | 5161.573 | 5158.252 |

Next, we generated a single self-regulating rAAV vector that would concurrently express a regulatory shRNA along with a transgene mRNA harboring a cognate 3’ UTR target site for the shRNA. To achieve this, we embedded the miR-30E-or miR-33-scaffolded TTR shRNA cassette into a chimeric intron residing between a Gaussia luciferase (GLuc) transgene and liver-specific TBG promoter and inserted a single fully matched binding site for the TTR shRNA within the 3’ UTR of GLuc **(Figure 1B and Supplementary Table 1).** This configuration allows expression of the regulatory shRNA and transgene mRNA to be coupled within a single transcriptional unit but permits each to be processed independently following precursor-mRNA splicing in the nucleus. We anticipated that once the shRNA is processed intracellularly, the resulting RISC-loaded antisense strand can mediate binding and cleavage of the rAAV-expressed transgene, which can then be reversed by exogenous treatment with REVERSIR **(Figure 1A).** *In vitro* evaluation of putative self-silencing rAAV plasmids demonstrated robust basal downregulation of the secreted GLuc reporter transgene with both miR-30E and miR-33 pri-miRNA scaffolds specifically when both a TTR shRNA and its complementary target site (TTR-ts) were present in the vector, relative to matched constructs expressing a NT shRNA or NT target site (NT-ts) (\**p*<0.0001 NT vs. TTR shRNA) **(Figure 1C and Supplementary Fig. 1C).** Given the need for intracellular processing of intronic shRNA into active RNAi effectors, longitudinal monitoring of transgene expression *in vitro* showed delayed suppression of GLuc transgene expression. Reporter levels were comparable for both self- and non-self-regulating designs at early timepoints with significant silencing only evident beginning at 14h post-transfection (shTTR/NT-ts vs. shTTR/TTR-ts 14h, 22h, 46h *p*<0.01) **(Figure 1C).** Co-transfection of 22-mer REVERSIR directed against the TTR shRNA resulted in recovery of GLuc protein and mRNA expression (TTR shRNA + 40nM TTR REVERSIR vs. TTR shRNA + 40nM NT REVERSIR \**p*<0.01). Consistent with a sequence-specific mechanism, we did not observe reversal of shRNA-induced knockdown of GLuc with NT REVERSIR **(Figure 1D and Supplementary Fig. 1D).**

We next assessed *in vivo* feasibility of this strategy using vectors expressing miR-33-embedded shRNA cassettes since prior studies had shown that this pri-miRNA backbone retained high efficacy without compromising rAAV genome integrity during viral replication.^25, 26^ We compared rAAV-GLuc reporter vectors expressing intronically-encoded miR-33-embedded TTR shRNA with either a fully complementary (TTR-ts) or scrambled (NT-ts) target site within the GLuc 3’UTR. Importantly, in line with our *in vitro* findings, expression of the self-silencing rAAV containing both an intronic miR-33 TTR shRNA and its complementary 3’ UTR binding site (shTTR/TTR-ts) led to a sustained knockdown (~76%) of secreted GLuc reporter signal, relative to the non-self-silencing (shTTR/NT-ts) vector design **(Figure 1E)**. GLuc serum levels recovered and were comparable to the control non-self-silencing vector within 7 days following molar equivalent subcutaneous (SC) dosing of either a short 9-mer [5 locked nucleic acid (LNA), 5 phosphorothioate (PS)] or long 22-mer (5 LNA, full PS) TTR REVERSIR, with induction lasting for at least 6 weeks (D49 \**p*>0.9 shTTR/NT-ts + vehicle vs. shTTR/TTR-ts + 9-mer TTR REVERSIR; *p*>0.9 shTTR/NT-ts + vehicle vs. shTTR/TTR-ts + 22-mer TTR REVERSIR). No significant GLuc induction was observed upon treatment with equivalent doses of the control NT REVERSIRs [Two-way ANOVA main effect of treatment F(1,160)=66.42 p<0.001; Tukey post-hoc p<0.0001 shTTR/NT-ts + vehicle vs. shTTR/TTR-ts + 22-mer or 9-mer NT REVERSIR). Consistently, qRT-PCR analyses also revealed rescue of *Gluc* transcript levels in the presence of TTR-REVERSIR relative to the NT REVERSIR control **(Figure 1F)**. These data successfully establish constitutive *in-cis* knockdown of transgene expression with intron-embedded shRNA and subsequent blockade of active RISC with REVERSIR.

While long-term induction may be desirable in certain contexts, the flexibility and utility of an RNAi/REVERSIR rheostat approach would be greatly enhanced by the ability to modulate the duration of REVERSIR action to achieve more transient upregulation of transgene expression. We hypothesized that reducing the metabolic stability of the REVERSIR may lead to its more rapid intracellular clearance, thereby facilitating resumption of RISC-mediated target engagement and silencing. Towards this end, we dosed mice transduced with self-silencing shTTR/TTR-ts rAAV with a 9-mer TTR REVERSIR designed with lower LNA and PS content (TTR REVERSIR 2 with 3 LNA and 3 PS) to diminish metabolic stability.^36, 37^ Both the evaluated doses (0.1 and 0.3 mg/kg) of TTR REVERSIR 2, led to initial recovery of GLuc transgene expression back to control levels by D6 (\**p*>0.1 shTTR/NT-ts + vehicle vs. shTTR/TTR-ts + TTR REVERSIR 2). Consistent with reduced metabolic stability and rapid clearance, dose-dependent resumption of GLuc silencing was observed over time, with levels reverting to those of NT REVERSIR-treated mice by D19 for the 0.1mg/kg (\**p*>0.3 TTR REVERSIR 2 vs. NT REVERSIR) and D47 for the 0.3mg/kg (\**p*>0.9) treatment groups **(Figure 1G).** A second peak in GLuc levels was observed following a re-challenge with another dose of TTR REVERSIR 2 on D47, highlighting the potential for repeated induction of transgene expression with tunable REVERSIR designs.

To demonstrate that this system can be used to control physiologically-relevant transgenes, we designed a TTR shRNA-embedded self-regulating AAV vector encoding mouse erythropoietin (*mEpo*) **(Figure 1H)**. Even though rAAV was administered at a high dose of 2 × 10^11^ GC, we observed >92% suppression of serum mEPO levels in mice injected with rAAV expressing TTR shRNA along with its cognate binding site relative to those harboring a non-targeted binding site. A single administration of a metabolically stable TTR REVERSIR but not NT REVERSIR resulted in complete recovery of mEPO expression that endured for at least 35 days **(Figure 1I).**

In contrast, treatment with a less stable REVERSIR design led to transient and dose-dependent increases in mEPO concentrations **(Figure 1J).**

Results from these studies suggest a viable single-agent strategy involving constitutive self-suppression of the rAAV transgene that may be counteracted by exogenous administration of REVERSIR to induce expression transiently or for a prolonged period of time.

### Transgene induction by REVERSIR blockade of exogenous siRNA activity

Extensive chemical modification of siRNAs, along with conjugation to novel ligands, have facilitated marked enhancements in potency, duration of action, and functional delivery to a broader range of extrahepatic tissues.^9–11,15^ Current siRNAs featuring enhanced stabilization chemistry (ESC) designs display durable target silencing that can persist for months in preclinical species and humans following a single administration.^9–11^ Accordingly, we postulated that the long-acting and reversible properties of siRNA activity may be uniquely suited to be repurposed as a programmable and user-specified system to achieve exogenous dosage control of rAAV transgene expression.

Using hepatotropic rAAV8 as a model, we sought to probe whether exogenous administration of GalNAc-conjugated siRNA and REVERSIR may enable on-demand modulation of AAV-expressed transgenes **(Figure 2A)**. Mice were injected with an rAAV8 vector encoding a bicistronic transcript composed of peripheral myelin protein-22 (PMP-22) and a GLuc reporter, with a single fully complementary binding site for a TTR-targeting siRNA inserted directly following the termination codon within the 3’ UTR **(Supplementary Table 1 and Figure 2B; *left***). Subcutaneous (SC) treatment with a single dose of a GalNAc-TTR siRNA resulted in sustained dose-dependent reduction of rAAV-driven serum GLuc expression, with the highest dose of 9mg/kg showing 71% silencing at 14 days and greater than 50% knockdown up to 45 days post-dose **(Table 2 and Supplementary Fig. 2A)**. Two weeks following siRNA treatment, mice received a single molar equivalent dose of either full-length (22-mer) or short 9-mer TTR REVERSIR, compared to chemistry- and length-matched NT REVERSIR as controls for sequence specificity **(Figure 2B; *right*).** Both the 22-mer and 9-mer TTR REVERSIRs efficiently counteracted silencing activity by the 9mg/kg siRNA dose, leading to prolonged and comparable induction of GLuc expression to levels similar to that of vehicle-treated controls for at least 4 weeks (D43 \**p*>0.1 PBS vs. siRNA + 9-mer or 22-mer REVERSIR; \**p*=0.7 22-mer TTR REVERSIR vs. 9-mer REVERSIR) **(Figure 2C and Supplementary Figure 2B).** In contrast, GLuc reporter knockdown was unaffected in the presence of NT REVERSIRs, with reporter suppression remaining comparable to mice treated with siRNA alone for the evaluated duration (D42 \**p*>0.9 siRNA + Vehicle vs. siRNA + 22-mer or 9-mer NT REVERSIR). Downstream qRT-PCR analyses showed that treatment with GalNAc-TTR siRNA resulted in significant reductions in mRNA levels of *Gluc*, which were restored to baseline following administration of 9-mer TTR REVERSIR but not the control NT REVERSIR **(Figure 2D).** To additionally illustrate that induction may be titrated to achieve varying levels of protein expression, we monitored GLuc expression following dosing of siRNA-treated mice with increasing amounts of 9-mer TTR REVERSIR. We observed dose-dependent upregulation of GLuc levels one week post-REVERSIR dosing, suggesting that REVERSIR can allow fine-tuning of AAV transgene expression to the desired level by modulating the magnitude of siRNA-mediated silencing activity **(Figure 2E).**

**FIGURE 2:**
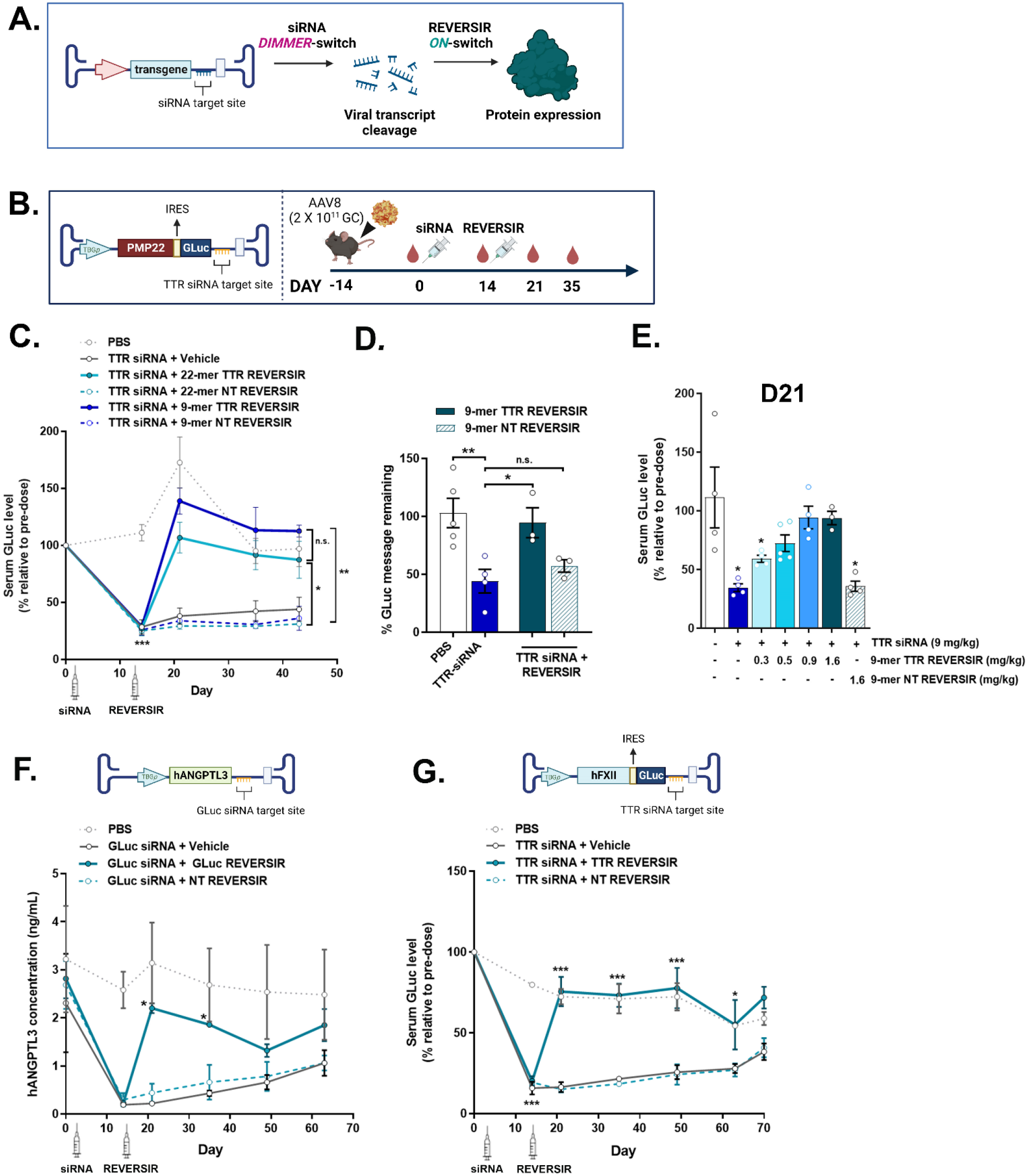
*In vivo* regulation of an AAV-delivered reporter transgene by exogenous delivery of siRNA and cognate REVERSIR. **(A)** Diagram depicting dual agent approach for rAAV transgene modulation. siRNA administration facilitates dampening of rAAV gene expression. Sequence-specific abrogation of siRNA activity with REVERSIR results in de-repression of viral mRNA transcripts and consequently increases therapeutic protein expression. **(B)** Schematic of viral vector encoding bicistronic expression of PMP-22 and GLuc, with TTR siRNA binding site in 3’ UTR **(left)** and experimental design **(right).** C57BL/6 mice were intravenously injected with 2×10^11^ GC of rAAV. Two weeks after rAAV administration, mice were injected SC with either vehicle or TTR siRNA at 9mg/kg (D0). This was followed two weeks later with a single molar equivalent dose of full-length 22-mer (3mg/kg) or 9-mer TTR REVERSIR (1.6mg/kg) and compared to vehicle or length-matched NT REVERSIR (D14) as controls. Blood was collected as indicated and terminal liver tissue harvested at D42. **(right) (C)** Quantification of serum GLuc levels at indicated timepoints normalized to pre-dose for each animal. **(D)** *Gluc* transcript levels in terminal liver tissue at D42 normalized to *Gapdh* control and plotted relative to PBS condition set to 100%. **(E)** Serum GLuc levels at D21 in mice transduced with 2 × 10^11^ GC of rAAV shown in (B) and treated with TTR siRNA (9mg/kg; D0), followed by varying doses of 9-mer TTR REVERSIR or NT REVERSIR (D14) at high dose alone as control. **(F)** Viral genome diagram for GLuc siRNA-mediated regulation of rAAV encoding human Angiopoietin-like 3 (hANGPTL3) transgene **(top).** Mice injected with 1.5 x 10^11^ GCs of rAAV were treated with 9mg/kg GLuc siRNA on D0, after which 1.6mg/kg 9-mer GLuc REVERSIR or NT REVERSIR was administered on D14. Bleeds were performed on D0, D14, D21, D35, D49, and D63 and plasma hANGPTL3 concentrations assessed by ELISA **(bottom). (G)** Viral genome schematic for TTR siRNA-mediated regulation of human Factor XII (hF12)-GLuc transgene **(top)**. Mice transduced with 2 × 10^11^ GC of AAV were treated with 9mg/kg TTR siRNA on D0, then after 2 weeks given 1.6mg/kg TTR REVERSIR or NT REVERSIR (D14). Serum GLuc intensity was measured on D0, D14, D21, D35, D49, D63, and D70, and plotted relative to pre-treatment with siRNA on D0 **(bottom).** Error bars represent s.e.m.

**Table 2:**
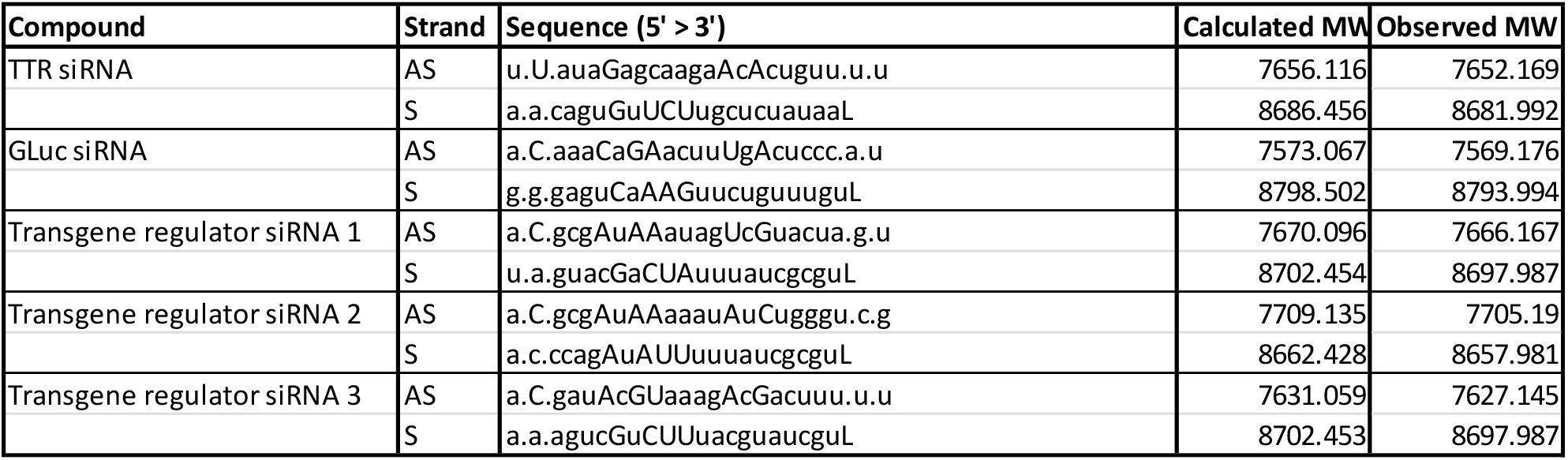
GalNAc-siRNA conjugates used in this study. Legend for chemical modifications is as follows: lower case nucleotides - 2’-*O*-methyl (OMe); upper case nucleotides – 2’-deoxy-2’-fluoro (F);. – phosphorothioate linkage; L – tri-*N*-acetylgalactosamine.

To demonstrate that exogenous dosing of siRNA with REVERSIR represents a generalizable approach for off- and on-state control of AAV transgenes, we evaluated two additional AAV vectors in which we varied either the identity of the expressed transcript and/or the regulatory siRNA sequence. Mice were transduced with an rAAV8 vector conferring expression of human Angiopoietin-like 3 (hANGPTL3) with a unique 3’ UTR target site for a GLuc siRNA **(Table 2)** to selectively target the exogenous transgene. Inherent potency of the GLuc siRNA was assessed *in vitro* using an on-target dual luciferase vector, demonstrating dose-dependent activity repression with an EC50 of 1.9nM **(Supplementary Fig. 2C)**. Co-transfection of GLuc siRNA with its cognate 22-mer REVERSIR yielded concentration-dependent recovery of luciferase expression to baseline levels, whereas the 9-mer GLuc REVERSIR displayed lower potency, recapitulating prior studies showing poorer performance of shorter REVERSIRs when delivered by lipid-based transfection **(Supplementary Table 1 and Supplementary Fig. 2D**). Treatment of AAV-transduced mice with 9mg/kg GLuc siRNA resulted in 91.7% suppression of serum hANGPTL3 protein concentrations at nadir (D14), with significant knockdown >80% detected up to D35 (D14, D21, D35 \**p*<0.05 PBS vs. siRNA + Vehicle). Subcutaneous administration of 3mg/kg 9-mer GLuc REVERSIR, but not NT REVERSIR, reverted serum hANGPTL3 to levels indistinguishable from vehicle controls with protein upregulation lasting for 3 weeks post-induction (D35 \**p*=0.7 PBS vs. siRNA + Gluc REVERSIR; \**p*<0.01 PBS vs. siRNA + 9-mer NT REVERSIR) (**Figure 2F and Supplementary Fig. 2E**). We also tested an rAAV construct encoding bicistronic expression of GLuc reporter with a 3’ UTR TTR siRNA target site as described previously. However, in this instance, we replaced the coding sequence of PMP-22 with that of a different gene, human Factor XII (hFXII). Durable and significant reduction in GLuc levels of >60% were detected up to D49, with maximal knockdown of 73.7% occurring at 2-weeks post-siRNA dose (D14 through 49 \**p*<=0.05 PBS vs. siRNA + Vehicle). In comparison to treatment with siRNA alone or with a NT REVERSIR, mice that received the TTR REVERSIR displayed significant upregulation of reporter expression up to D63 (D21 through D63 \**p*<0.01 siRNA alone vs. siRNA + TTR REVERSIR; \**p*<0.01 TTR REVERSIR vs. NT REVERSIR) (**Figure 2G and Supplementary Fig. 2F**). These data highlight the generalizable nature of this regulatory switch approach for off- and on-state control of various AAV transgenes.

Collectively, these results indicate that exogenous delivery of GalNAc-conjugated siRNAs and REVERSIR represents a generalizable strategy for titratable and on-demand control of transgene expression following rAAV transduction.

### Identification of potent and specific transgene regulator siRNAs

Practical *in vivo* application of an RNAi-based regulatory switch for AAV vectors necessitates highly selective and unique siRNA sequences that allow RNAi activity to be exclusively directed towards the intended exogenously-expressed transgene, while simultaneously mitigating potential sequence-dependent off-target effects on the endogenous transcriptome. Using bioinformatic approaches, we identified three candidate AAV transgene regulator siRNAs (TR-siRNAs) that were predicted to have favorable thermodynamic properties for RISC loading and RNAi functionality. Importantly, all the siRNAs were designed to have little to no sequence complementarity to any annotated transcripts in human, cynomolgus monkey, rat, and mouse **(Supplementary Table 1).** To probe their intrinsic silencing efficacy, we assessed reporter activity *in vitro* in response to increasing doses of the TR-siRNAs in the presence of dual luciferase vectors bearing a single copy of a perfectly complementary target site in the 3’ UTR. All three siRNAs exhibited significant dose-dependent on-target repression of luciferase reporter activity, with IC_50_ values in the low nanomolar range [0.18nM, 0.71nM, and 0.44nM for siRNAs 1-3 respectively) **(Figure 3A)**. To evaluate the potential for microRNA-like off-target knockdown, we co-transfected the siRNAs with luciferase sensors containing either a single copy or four tandem repeats of a site complementary to the seed region (nucleotides 2-8) of the siRNA antisense strand. No significant microRNA-like off-target luciferase reporter repression was observed at doses ranging from 0.64pM to 50nM for siRNAs 1 and 2. siRNA 3 showed significant knockdown at doses ≥10nM with the off-target reporter containing 4 sites, but no off-target activity was observed at any dose for the reporter harboring a single seed-matched site (**Figure 3A**).

**FIGURE 3:**
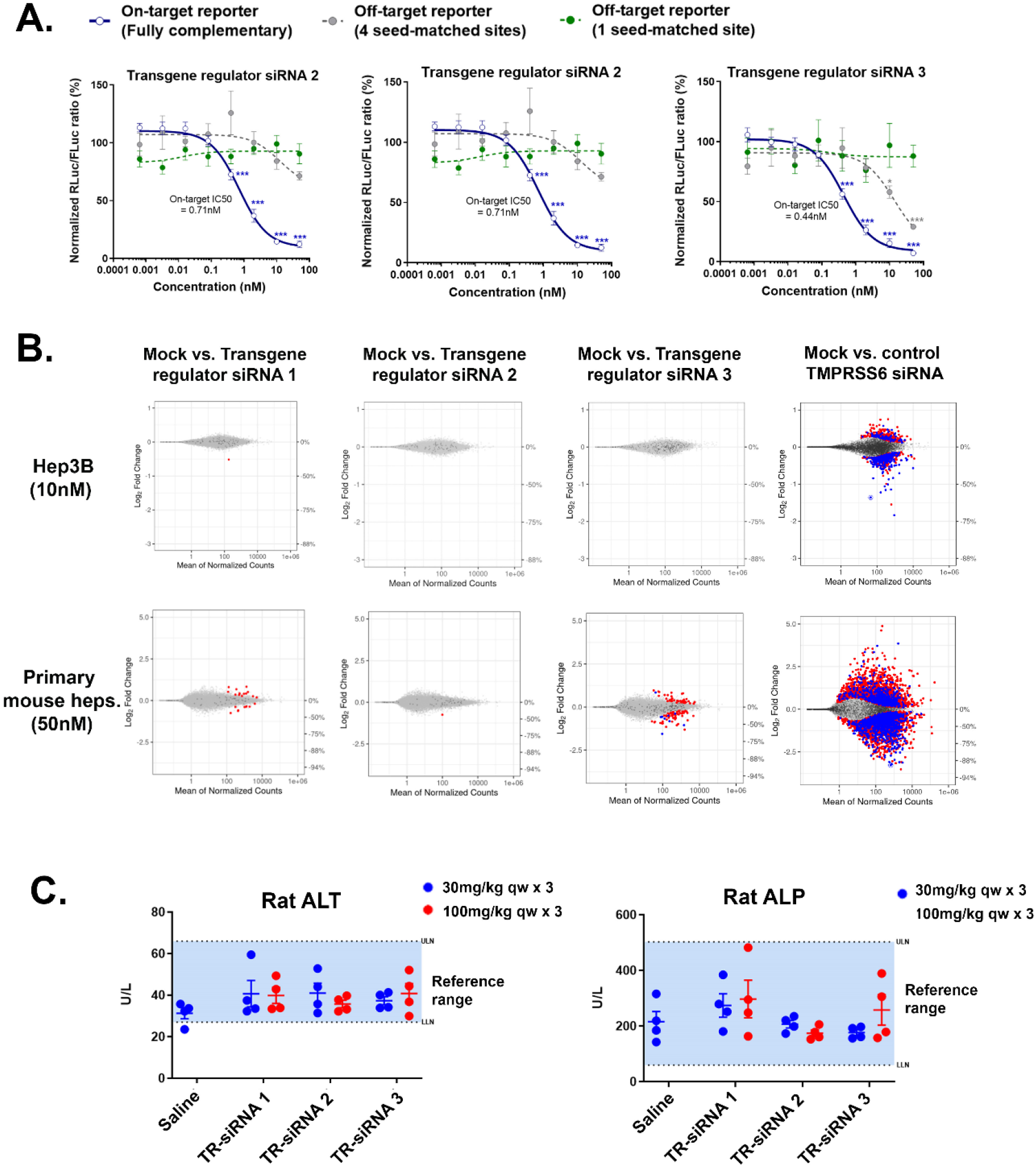
*In vitro* characterization of on- and off-target activity of transgene regulator siRNA sequences. **(A)** On-target silencing efficacy (solid blue line) of three lead transgene regulator siRNA (TR-siRNA) sequences as assayed by co-transfection of serially titrated doses of siRNA with dual luciferase sensors containing a perfectly matched binding site in the 3’ UTR of Renilla luciferase (RLuc). Seed-mediated off-target repression was similarly assessed by dose-response activity of siRNA in the presence of luciferase reporters bearing either 1 (green dashed line) or 4 tandem (grey dashed line) seed-matched target sites. RLuc/FLuc ratios were normalized to the mock-transfected control (no siRNA) condition set at 100% and plotted as mean of 3-6 replicates ± s.e.m. **(B)** MA plots depicting differential gene expression analysis of RNA-seq data obtained from transfection of TR-siRNAs and a *Tmprss6*-targeting siRNA control in Hep3B cells (top; 10nM dose harvested at 24h) and primary mouse hepatocytes (bottom; 50nM dose harvested at 48h). Dots represent individual mRNAs, average normalized read counts across replicates, and log2 fold change relative to mock transfected controls. ‘Red’colored dots represent genes with significant differential expression (false discovery rate <0.05) but no canonical seed-matched sites within their 3’ UTRs (8-mer, 7mer-m8, and 7mer-A1). ‘Blue’-colored dots indicate genes with significant differential expression and canonical seed-matched sites within their 3’ UTRs. ‘Dark gray’ dots denote genes that contain canonical 3’ UTR seed-matched binding sites but are not significantly differentially expressed (*N* = 4 technical replicates). The circled dot represents on-target knockdown. **(C and D)** Rat toxicity evaluations of TR-siRNAs **(C)** Serum alanine aminotransferase (ALT) and alkaline phosphatase (ALP) at necropsy (24 hours after last dose of siRNA). N = 4 males (6-8 weeks old) per group; qw weekly dosing. Error bars represent s.e.m.

To characterize the potential for sequence-dependent and -independent off-target effects more broadly, we transfected these candidate TR-siRNAs into Hep3B cells and primary murine hepatocytes to assess their global impact on the human and mouse transcriptomes by RNA-seq. As designed, there were relatively few transcripts that contained canonical seed matches (8mer, 7mer-m8, and 7mer-A1) to the TR-siRNA antisense strands within their 3’UTRs. Across both cell types, all three TR-siRNAs generally displayed minimal to no transcriptional dysregulation when comparing log2 fold-changes in mRNA expression levels relative to mock control for genes harboring canonical seed-matched (blue dots) sites as well as those without canonical seed matches (red dots). Additionally, 2 of the 3 siRNAs showing no transcripts with a magnitude of dysregulation exceeding 2-fold **(Figure 3B and Supplementary Table 2).** A *Tmprss6*-targeting siRNA that was assayed in parallel showed robust on-target transcript knockdown, confirming successful transfection of all the evaluated siRNAs.

To confirm that the observed lack of off-target gene dysregulation translates to a favorable safety profile *in vivo*, we administered the three TR-siRNAs at toxicological doses in rats. Rats received once weekly subcutaneous injections (qw x 3 doses) of 30 or 100mg/kg siRNA, representing 2-3 log increase from the pharmacological dose range. All three siRNAs showed no significant liver enzyme elevations **(Figure 3C and Supplementary Fig. 3A).** RNA-seq evaluation of rat livers subjected to the toxicological dose regimen showed a dose-dependent increase in differentially expressed gene transcripts, particularly amongst those lacking any canonical seed-matched binding sites. Changes in the expression of these transcripts could reflect indirect secondary effects or other non-canonical matches to the TR-siRNA. For instance, these could include sites with weaker complementarity to the siRNA seed region (6mer, etc.) or complementarity to the 3’ supplemental region of the antisense strand. Nonetheless, these siRNA sequences exhibited no evidence of transcriptional dysregulation among canonical seed-matched transcripts, indicating a substantial lack of microRNA-like off-target gene signatures at toxicological doses **(Supplementary Fig. 3B).**

Overall results from our *in vitro* RNA-seq and rat toxicity screening efforts establish these siRNAs as potent RNAi triggers with high on-target specificity. These TR-siRNA sequences display minimal propensity for off-target gene disruption and *in vivo* toxicity, lending initial support for their use as regulators of exogenous transgenes with potentially favorable safety profiles at least in the context of liver-directed applications.

## DISCUSSION

A lack of broadly applicable methods to control the dosage of therapeutic transgenes is a central challenge surrounding rAAV gene therapies, since most currently rely on persistent constitutive expression following a single administration. The ability to refine transgene dosage to within the targeted therapeutic range or suppress expression in case of adverse events represent important features that could maximize the safety of rAAV-based therapies. RNA-based molecular switches are emerging as attractive candidates to regulate transgene expression from AAVs owing to their small size, sequence specificity, and reduced potential for immunogenicity.^19^ Recent developments in riboswitches based on steric oligo blockade of self-cleaving ribozyme activity and small molecule regulation of alternative RNA splicing have shown promise in preclinical models but their applicability in clinical settings is unknown.^38, 39^ We report a generalizable and clinically viable approach for dosage control of rAAV-delivered cargos involving dynamic, robust, and reversible control via RNAi with benefit of infrequent dosing. We demonstrate reversal of RNAi-mediated transgene knockdown with REVERSIR in both a dual agent setting involving exogenous administration of chemically modified siRNA, and a single agent setting where co-delivery of vectorized shRNA enables constitutive silencing. Both approaches provide durable dampening of transgene expression that may be abrogated with REVERSIR to yield weeks- to months-long elevation in protein levels in mice following a single dose of the inducer.

Unlike riboswitch designs that depend on self-cleaving ribozyme activity or splicing-dependent transgene cassettes, both of which afford extremely tight control in the off-state, an RNAi mechanism is unlikely to completely abolish basal transgene expression. Nevertheless, clinical trials evaluating investigational RNAi therapeutics have shown robust and prolonged target knockdown; results from the ORION Phase 3 study of a GalNAc-siRNA targeting PCSK9 support a once-every-6-month dosing regimen.^8^ We showed dose-dependent lowering of rAAV-expressed protein levels that were maintained for over a month in rodents after a single dose of siRNA or were constitutively suppressed in the case of vectorized shRNA.

Despite the slower kinetics of onset and inability to wholly inactivate transgene expression, there are certain settings where the durability of RNAi activity and the potential for infrequent dosing could have distinct utility for the regulation of rAAV gene therapies. For instance, RNAi may be well suited for regulation of certain cargoes, such as monoclonal antibodies, where fine tuning antibody concentrations below the required threshold for therapeutic effect may be sufficient to minimize adverse effects.^40^ One could also consider using RNAi to address the challenge of dose scaling and management for highly active transgenes, such as the Padua variant of FIX, where small differences in vector dose could lead to exaggerated changes in protein production.^6^ Systemic administration can also produce variable levels of transgene expression in response to the same vector dose, consequently increasing the risk of adverse toxicity if protein expression far exceeds the therapeutic range. This was recently highlighted in trials evaluating rAAV-FVIII for the treatment of hemophilia A where a high degree of variation in FVIII activity levels was observed among participants.^2, 41^ One could envision using an RNAi-based approach to titrate transgene expression to within the therapeutic window and thereby mitigate potential negative consequences associated with over-production of the therapeutic protein, such as increased thrombotic risk in the case of high FVIII levels. In addition to initial dose titration after rAAV dosing, RNAi-based approaches could enable transgene expression to be dynamically modified as the disease state evolves. Incorporation of an RNAi-based safety switch could allow temporary cessation of treatment if needed. With vectorized shRNA co-expression, the potential to delay stable transgene expression until after immune responses to rAAV vector subside represents another potential key benefit.^42^ Interestingly, with vectorized shRNA co-expression, the transient expression of transgene is seen before the RNAi effect is observed. This can be attributed to delayed kinetics of shRNA processing versus transgene mRNA translation and as such could find utility in approaches where only transient expression is needed upon rAAV dosing.

A key piece of data that emerged from our studies was that REVERSIR treatment afforded complete recovery of the full range of transgene expression that was observed in the absence of the RNAi regulatory module. A previously reported RNAi on-switch design based on ligand-promoted occlusion of a microRNA target site with a competing complementary strand yielded a maximal recovery of ~20% of non-regulated expression, likely owing to the remarkable efficiency with which RISC recognizes and binds its targets.^23^ Our observations indicate that blocking RNAi activity directly at the level of catalytic RISC provides a significantly more efficient and robust means of regulating the on-state.

Whereas our approach facilitated maximal recovery of transgene expression, the overall dynamic range for induction was still relatively modest, with ~5-10 fold regulation of reporter gene expression detected *in vivo*. However, in line with previous studies showing that more efficient RNAi-mediated silencing results in wider dynamic ranges for induction, we observed the most significant fold upregulation with rAAV-expressed hANGPTL3 where we were able to achieve >90% suppression of basal transgene expression with siRNA.^23^ On the other hand, on-state control via regulation of alternative splicing with small molecules or ribozyme activity with steric oligos achieve a much larger magnitude (>100-fold) but a relatively transient duration of transgene induction.^38, 39^ As a result, chronic administration of the inducer might be required to maintain persistent expression of the therapeutic transgene, particularly for proteins and peptides with a short half-life. In line with the durability of RNAi responses, we showed that a single dose of REVERSIR produced weeks-long elevations in GLuc reporter, which has an extremely short serum half-life of 20 minutes. We also demonstrated that duration of induction can be controlled by modifying the metabolic stability of the REVERSIR, thereby adding a layer of versatility for different gene therapy applications. Additional refinements to the metabolic stability and dose of the REVERSIR could feasibly shorten timescales of induction even further.

We also report potent siRNA sequences with high specificity that can be incorporated into episomal vectors to selectively regulate exogenous transgene expression. Several studies have shown that hybridization of the siRNA seed region to complementary sequences within the 3’ UTR of mature transcripts is the principal driver of RNAi-mediated off-target gene dysregulation and *in vivo* toxicity.^43, 44^ Therefore, we prioritized siRNAs with seed-matched target sequences that occur at low frequency and whose full-length sequences lack homology to expressed transcripts across mouse, rat, cynomolgus monkey, and human genomes. We saw minimal to no seed- or non-seed-mediated dysregulation of endogenous mRNA expression in both human and mouse hepatic cell lines at high doses. All three TR-siRNAs demonstrated a favorable safety profile in a repeat-dose rat toxicity study. These data suggest that these siRNAs may be administered at relatively high doses if needed with minimal risk of the off-target effects. Overall, these features increase their potential for clinical translation with favorable safety profile. Our current studies utilized a regulatory element consisting of a single fully-matched binding site within the 3’ UTR of the rAAV transgene. The sensitivity of the RNAi-driven regulation could be further influenced by modifying the local sequence around the target site or altering the proximity of the target site to the transgene stop codon.^45^

While our current investigations are limited to control of hepatocyte-directed transgene expression with GalNAc-conjugated siRNAs and REVERSIRs, novel delivery solutions such as C16 may broaden the scope of RNAi-based strategies for control of AAV vectors targeting a wide range of tissues.^15^ For extrahepatic applications, constitutive basal transgene silencing via vectorized RNAi delivery might be preferable since it obviates the need for exogenous siRNA and enables a single agent strategy involving dosing of REVERSIR alone.

In summary, our data suggest that an RNAi-based rheostat could be a potent and clinically adaptable tool for expression control from rAAV vectors, with a unique profile and use case compared to other transgene regulatory modalities.

## MATERIALS AND METHODS

### Plasmids

On- and off-target reporters were generated by sub-cloning a DNA fragment containing a fully complementary (23-mer) or partial siRNA target sites into the psiCHECK2 vector (Promega C8021) between *Xho1* and *Not1* restriction sites. The on-target reporter plasmids contain a single site with full complementarity to the antisense strand cloned into the 3’-UTR of Renilla luciferase. Off-target reporter plasmids contain either one or four tandem seed-matched sites, complementary to antisense positions 2 to 9, separated by a 19-nucleotide spacer (5’-TAATATTACATAAATAAAA-3’) cloned into the 3’-UTR of Renilla luciferase.^46, 47^

rAAV vectors were generated using Vector Builder’s or Blue Heron’s single-stranded AAV vector backbone with a TBG promoter, KOZAK sequence preceding the transgene, and bGH-poly A signal. For vectors harboring pri-miRNA-adapted shRNAs, the chimeric intron cassette from *pAAVsc-CB-PI-GLuc* was sub-cloned between the promoter and transgene elements, and the shRNA fragment was cloned into the PpuM1 site. The miR-33 and miR-30E shRNA sequences were designed as described previously.^25, 27^ A perfectly complementary siRNA target site was inserted immediately downstream of the GLuc and EPO transgenes, without any intervening spacer sequence. rAAV vectors used in Figure 2 expressed a mono- or bicistronic transcript with a fully complementary siRNA target site after the transgene stop codon, separated by a NotI restriction site. Luciferase reporter and vectorized shRNA insert sequences are detailed in **Supplementary Table 1**.

### Care and use of laboratory animals

All procedures and protocols performed on rodents were approved by Alnylam’s Institutional Animal Care and Use Committee (IACUC) and were compliant with guidelines set by local, state, and federal regulations. Female C57BL/J mice and male Sprague Dawley rats between 6-8 weeks of age were obtained from Charles River Laboratories and allowed to acclimate in-house for 48 hours prior to initiation of studies. Adult mice were group housed (up to 5 sex-matched animals per cage) on a standard 12:12 hour light-dark cycle and provided access to food and water *ad libitum*.

### AAV injections, serum and terminal tissue collections in mice

Single-stranded rAAV8 vectors were generated, purified, and titered either at University of Massachusetts Viral Vector Core or Signagen Laboratories. Viral stocks were diluted in sterile 1X PBS,and administered intravenously by tail vein injection at the indicated titer in a total volume of 100μL (amount of virus injected is reported in the figure legends). Two weeks after AAV administration, mice were subcutaneously injected with GalNAc-conjugated siRNA or GalNAc-conjugated REVERSIR, or phosphate buffered saline (PBS) as control, in a total dosing volume of 10μL/g. All siRNA and REVERSIR test articles were diluted to the appropriate dosing concentration in 1X PBS.

Blood was collected by alternating retro-orbital bleeding under isoflurane anesthesia in accordance with IACUC approved protocols. For serum samples, blood was collected in Becton Dickinson serum separator tubes (Fisher Scientific, BD365967), kept at room temperature for 1 h and then spun in a micro-centrifuge at 21 000 × *g* at room temperature for 10 min. For plasma samples, blood was collected in Becton Dickinson plasma (K2EDTA) separator tubes (Fisher Scientific, BD365974), kept on ice for 30 min before being centrifuged at 10 000 × *g* at 4°C for 10 min. Both serum and plasma samples were aliquoted and transferred to 96-well plates for storage at −80°C. Animals were sacrificed at the indicated days, after which terminal livers were harvested, flash frozen in liquid nitrogen, and stored at −80C until further downstream analysis.

### Rat toxicity studies

Test articles were diluted with 0.9% NaCl to achieve appropriate dosing concentrations and dosed subcutaneously on the upper back to male Sprague Dawley rats in a dose volume of 5 mL/kg with N = 4 animals/group. Rats were sacrificed and livers and blood collected on Day 16. Randomization was performed using the partitioning algorithm in the Pristima^®^ Suite (Xybion) that avoids group mean body weight bias. Investigators were not blinded to the group allocation during the experiment or when assessing the outcome. Whole-venous blood was collected into serum separator tubes (BD Microtainer) and allowed to clot at room temperature for 30 min prior to centrifugation at 3000 RPM (1489 × g) for 10 min at 4 °C. Serum was then aliquoted and stored at −80 °C until analyses. Serum chemistries were analyzed using the AU400 chemistry analyzer (Beckman Coulter-Brea, CA, USA), with reagents provided by Beckman Coulter, Randox, and Sekisui Diagnostics.

### Oligonucleotide synthesis

All oligonucleotides were synthesized on an MerMade 192 or MerMade 12 synthesizer according to previously published protocols.^15,30,48^ Sequences of all siRNAs and REVERSIRs used in these study are listed in **Supplementary Tables 1 and 2.**

### Bioinformatic prediction of transgene regulator siRNA sequences

Selection of the molecular switch duplexes was informed by conventions used for therapeutic siRNA development candidates by Alnylam: 21 nucleotide sense strand, and a 23 nucleotide antisense strand, forming a 2 base overlap at the 3’-end of the guide strand.

A set of all decamers was generated in silico to represent the first 10 bases of a candidate antisense (guide) sequence (1,048,576 sequences). miRNA seed sequences were retrieved from the miRbase database for *Homo sapiens, Mus musculus, Rattus norvegicus, Macaca mulatta* and *Macaca nemestris*. The non-human primate species were selected as proxies for *Macaca fascicularis* (cynomolgus monkey), which is not represented in miRbase. Decamers containing seed sequences (bases 2-7) matching the miRNA seed sequences were removed from further consideration (487,186 decamers remaining).

Each remaining decamer was annotated with a predicted quiescence score based on a proprietary off-target prediction algorithm. Those with scores in the lowest quartile (predicted most quiescent) were retained (155,374 decamers), and others removed from consideration.

The frequency of heptamers found in the human transcriptome (NCBI RefSeq), as was computed by aligning transcripts for each gene, and counting all heptamer substrings in the global alignment consensus and non-overlapping regions. Decamers were then annotated with the frequency of the heptamer in the seed (positions 2-8), and those in the lowest decile heptamer frequency were retained (1,767 decamers) and the remainder removed from consideration.

The remaining decamers were then aligned to the human, mouse, rat, and cynomolgus monkey transcriptomes using BLASTN with the parameters “-task blastn-short -dust no -evalue 1000 -ungapped - perc_identity 100” to count the number of occurrences of each decamer within each transcriptome. The decamers were then sorted in ascending order based on the total number of identical alignments on the reverse complement strand, then predicted quiescence, and number of seed matches in the human transcriptome.

For each decamer, 10,000 random 13-mer sequences were created and appended to create 10,000 candidate 23-mer siRNA guide sequences with a common 10-mer prefix. Each 23-mer was then aligned to the transcriptomes of human, mouse, rat, and cynomolgous monkey using a weighted ungapped alignment of the 23-mer to the transcripts, (mismatch penalty for positions 2-9 is 2.8, for positions 10-11, 1.2, for positions 12-19, 1.0, and for positions 1 and 20-23, 0.0). For each candidate decamer prefix, the 23-mers with the top 10 worst alignment profiles (most poorly aligned to the transcriptomes) were retained. Sequences were sorted by their alignment score and predicted quiescence, and top candidates were selected.

### ELISA assays

Circulating AAV-expressed human ANGPTL3 protein levels were measured from plasma using commercially-available ELISA kits (plasma diluted 1:4 and used with R&D Systems #DANL30). The assay is specific for detection of human ANGPTL3 protein, with no significant cross-reactivity to other related angiopoietin molecules or mouse ANGPTL3. Mouse EPO concentrations were measured with Mouse EPO Quantikine ELISA kit from R&D Systems MEP00B (serum diluted 1:150). TTR protein levels were measured with mouse prealbumin kit from ALPCO, 41-PALMS-E01 (serum diluted 1:4000). All assays were performed following the manufacturer’s protocols.

### Cell lines and transfection

Cos-7 (ATCC CRL-1651) and HepG2 (ATCC HB-8065) cells were grown in DMEM and EMEM, respectively, both supplemented with 10% heat-inactivated FBS and 1% glutamine and maintained in a humidified incubator at 37°C, 5% CO2. Plasmids and siRNAs were co-delivered by reverse transfection using Lipofectamine 2000 (Thermo Fisher Scientific 11668) for Cos-7 cells and Lipofectamine 3000 for HepG2 cells, following the manufacturer’s protocol.

### Luciferase reporter assays

#### siRNA on-target and off-target reporter evaluations

Cos7 cells were co-transfected with 5ng psiCHECK2 reporter plasmid and the specified amounts of siRNA duplexes (serially diluted in PBS) in a 384-well plate format at a density of 5×10^3^ cells per well. To assess REVERSIR-mediated rescue of knockdown by vector-encoded shRNAs expressed *in trans*, Cos7 cells were co-transfected in the same 384-well format with 70ng of shRNA expression plasmid, 5ng of psiCHECK2 reporter, in addition to indicated amounts of the specified REVERSIR molecules (serially diluted in PBS). 48 hours post-transfection, Firefly (transfection control) and Renilla (target) luciferase activities were sequentially measured using the Dual-Glo Luciferase Assay System (Promega E2920) and detected on a Spectramax M plate reader (Molecular Devices). The Renilla signal was normalized to Firefly signal for each well and expressed as a percentage relative to control wells transfected with reporter alone without siRNA or non-targeting shRNA plasmid. All transfections were performed at least in triplicate.

#### In vitro characterization of self-silencing AAV constructs

HepG2 cells were seeded at a density of 2×10^4^ cells per well in a 96-well plate and co-transfected with 16.6ng intronic shRNA-containing GLuc expression plasmid along and 20nM or 40nM REVERSIR. As an internal control to normalize for transfection efficiency, 3.3ng of PGK-driven Luc2 (pGL4.53[luc2/PGK]) vector was also co-transfected, constituting 17% of the total transfected DNA. Reverse transfections were carried out with 0.1 μL P3000 and 0.2μL Lipofectamine 3000 per well and allowed to proceed for 6 hours after which the media was replaced. Expression of GLuc and Luc2 reporters was measured 48 hours after transfection. To measure secreted GLuc levels, cell culture supernatant from each sample was diluted 1:50 in EMEM. 5μL of diluted supernatant and 50μL of assay buffer containing 3μM coelenterazine substrate (Selleck Chem S7777; stocks made up to 1mM in DMSO and subsequently diluted to 3μM in IX PBS) were transferred to each well of a white opaque 96-well plate and read on a Spectramax L microplate luminometer. Plate was dark-adapted to minimize auto-luminescence and injection speed was set to 250μL/s, followed by 2s shake and 1s signal integration time per well. To determine cellular Luc2 expression, cells in each well were first lysed with 50μL ice-cold 1X passive lysis buffer (Promega E1941), allowed to incubate on an orbital shaker for 10 minutes at room temperature, after which 50μL of ONE-Glo EX luciferase reagent (Promega E8110) was added. Luciferase signals were detected on a SpectraMax M plate reader 5 minutes later. Ratio of GLuc to FLuc intensity was computed for each well, and expressed as percent normalized to a matched NT shRNA-expressing condition.

#### Longitudinal monitoring of secreted GLuc levels in vivo

5μL of serum was assayed as described above on a Spectramax L luminometer. All serum samples corresponding to a study were run simultaneously under uniform conditions and resulting GLuc signals at an individual timepoint were expressed as percent relative to that at D0 (pre-treatment with siRNA or REVERSIR) for each animal.

### RNA isolation and RT-qPCR evaluations

96-well plates of cells were directly homogenized in 100μL RLT buffer (Qiagen) containing 10μL/mL beta-mercaptoethanol and allowed to incubate on an orbital shaker for 10 minutes at room temperature. RNA was extracted using the Qiagen RNeasy 96 kit (#74181), with additional incorporation of an on-column DNase digestion step (Qiagen 79254). For *in vivo* samples, powdered liver (~10 mg) was resuspended in 900 μL QIAzol (RNeasy 96 Universal Tissue Kit, Qiagen, 74881) and homogenized at 25/s for 1 min at 4°C using a TissueLyser II (Qiagen, 85300). Samples were incubated at room temperature for 5 min followed by addition of 180 μl chloroform. Samples were vigorously mixed, followed by a 10 min incubation at room temperature. Samples were spun at 12 000 × *g* for 15 min at 4°C, the supernatant was moved to a new tube, and 1.5 volumes of 70% ethanol was added. Samples were then purified using a RNeasy 96 Universal Tissue Kit (Qiagen, 74881) with on-column DNase digestion. RNA was eluted from the RNeasy spin columns with 50 μl RNAse-free water (Ambion) and quantified on a Nanodrop (Thermo Fisher Scientific). 0.5 μg of purified RNA was reverse transcribed using either the iScript gDNA Clear cDNA Synthesis Kit (BioRad, #1725034) or Maxima H Minus First Stand cDNA Synthesis Kit (Thermo Fisher Scientific M1681). RNA was DNase-treated prior to cDNA synthesis and oligo(dt)_18_ primers were use, where applicable. The product was diluted 1:2 in RNase-free water and subjected to quantitative real-time PCR (qRT-PCR) using gene-specific TaqMan assays (Thermo Fisher Scientific 4331182) for mouse TTR (Mm00443267_m1), human PMP22 (Hs00165556_m1), Luc2 (forward primer: TAAGGTGGTGGACTTGGACA, reverse primer: GTTGTTAACGTAGCCGCTCA, FAM-MGB probe: CGCGCTGGTTCACACCCAGT), and GLuc (Catalog #4441114 ARKA6G2). Levels of mouse (Mm99999915_g1) or human (Hs99999905_m1) GAPDH were used as endogenous normalization controls. Real-Time PCR was performed in a Roche LightCycler 480 using LightCycler 480 Probes Master Mix (Roche, 04707494001). No-reverse transcriptase enzyme controls were performed on a few samples in each study to ensure lack of viral or genomic DNA contamination. All RT-qPCR data were analyzed using the ΔΔCt method.

### RNA-seq methods

Primary Mouse Hepatocytes (BIOIVT, Cat # M005052-P, Lot GBW) were transfected in 384-well plates (5000 cells per well) with siRNA or DPBS (mock control) at a final concentration of 50 nM using Invitrogen Lipofectamine^®^ RNAiMAX (Invitrogen, Carlsbad, CA, Catalog No. 13778-150). After 48 hours, cell were lysed in lysis buffer (Tris HCl pH 7.5 100mM, LiCl 500mM, EDTA pH 8.0 10mM, LiDS 1%, DTT 5mM supplemented with TURBO™ DNase, Thermofisher #AM2238) for 30 min. Subsequently, whole transcriptome mRNA-enriched RNA-Seq libraries were constructed from cell lysates using a KAPA mRNA Capture Kit and mRNA HyperPrep Kit (Roche) as per the manufacturer’s protocol. RNA-Seq libraries were quantified by low depth sequencing on Illumina iSeq instrument. Equal amounts of each library/sample were pooled and sequenced on an Illumina NovaSeq instrument with 2×150bp paired-end settings, according to manufacturer’s instructions.

Raw RNAseq reads were filtered with minimal mean quality scores of 28 and minimal remaining length of 36, using the ea-utils software fastq-mcf v1.05.^49^ Filtered reads were aligned to the *Mus musculus* genome (GRCm39/mm39) using STAR (ultrafast universal RNAseq aligner) v2.7.9a.^50^ Uniquely aligned reads were counted by featureCounts v2.0.2 with the minimum mapping quality score set to 10.^51^ Differential gene expression analysis was performed by the R package DESeq2 v1.34.0 with the betaPrior parameter set to TRUE to shrink log2 fold-change estimates for noisy, low-count genes.^52^

### Statistical Analysis

Statistical analyses were performed using GraphPad Prism v.7 software. Data were analyzed by one-way or two-way analyses of variance (ANOVA) followed by,Tukey’s, Dunnett’s, or Holm-Sidak’s post hoc tests for multiple comparisons. In certain instances where indicated, individual unpaired two-tailed *t*-test or multiple *t*-testing (one per row) with Holm-Sidak correction was employed. All data are presented as mean +/- s.e.m. and differences between groups considered significant if *** *p*<0.0001, *** *p*<0.001, \*\**p*<0.01, or \**p*<0.05. Relevant non-significant differences between groups are highlighted with “n.s.”

## Data Availability

The raw RNA sequencing data reported in this manuscript have been deposited to the NCBI Gene Expression Omnibus and are accessbile through GEO Series accession number GSE214065.

## Acknowledgments

The authors thank Alnylam’s *in vivo* Sciences Team for conducting the safety studies and the Pathology team for generating the clinical pathology data. Schematics were created with BioRender.com.

## Author contributions

Conceived studies: M.S., J.M., C.K., M.M., K.F., V.J. Designed studies and experiments: M.S., J.M., I.Z., M.K.S., C.K., S.A., V.J. Performed experiments: M.S., J.M., C.K., T.N., S.A., T.R., M.A.A., K.W., Ty.C., T.C., E.S., S.S.M., A.B. Analyzed and interpreted data: M.S., K.B., D.B., J.B., S.A., J.M., I.Z., C.K., C.R.B., A.B.C., K.S., V.J. Supervised: M.M., K.F., V.J. Prepared the manuscript with input from all authors: M.S., V.J.

---

**SUPPLEMENTARY FIGURE 1:**
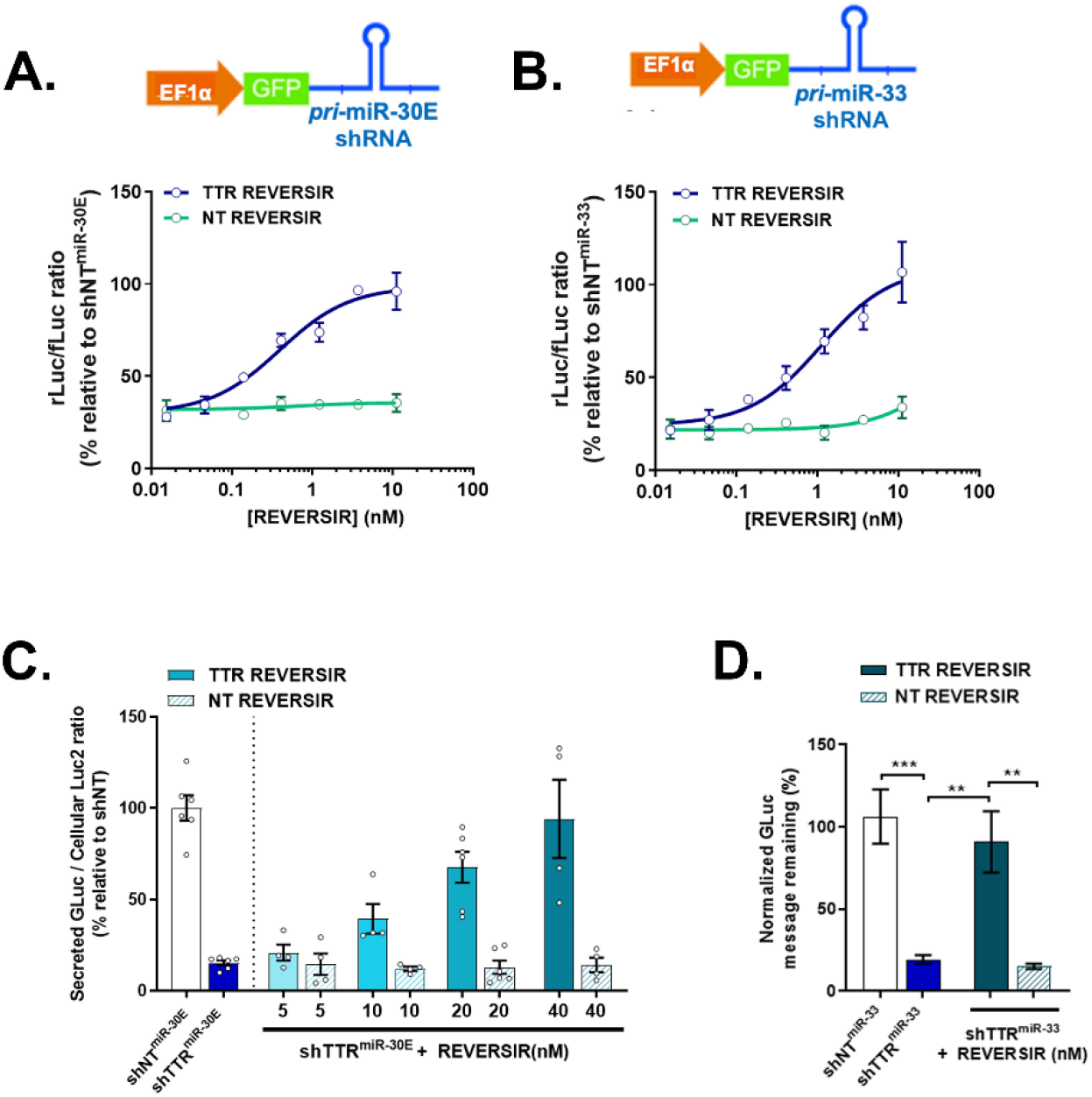
Additional *in vitro* and *in vivo* analyses supporting rAAV regulatory switch leveraging intronically-expressed shRNA and REVERSIR. **(A and B)** *In vitro* assessment of REVERSIR-mediated reversal of target silencing by miRNA-mediated shRNA in the dual luciferase reporter assay. Cos7 cells were co-transfected for 48 hours with luciferase reporter plasmid and GFP marker constructs expressing miR-30E- **(A)** or miR-33- **(B)** embedded TTR or NT shRNAs, along with increasing concentrations of 22-mer TTR or matched NT REVERSIR. **(C)** Validation of GLuc transgene silencing with intronic miR-30E-shRNA-containing **(right)** rAAV constructs and subsequent induction with increasing doses of REVERSIR in HepG2 cells. AAV constructs were co-transfected with FLuc control plasmids for normalization at 5:1 molar ratio. GLuc and FLuc intensities were assayed in cell culture supernatant and lysate, respectively, and GLuc/FLuc ratios expressed as % relative to shNT-expressing AAV plasmid. **(D)** Quantification of GLuc mRNA levels by qRT-PCR in HepG2 cells 48h following transfection with indicated rAAV plasmids and REVERSIR. GLuc transcript levels were normalized to FLuc mRNA as internal control. All error bars represent s.e.m.

**SUPPLEMENTARY FIGURE 2:**
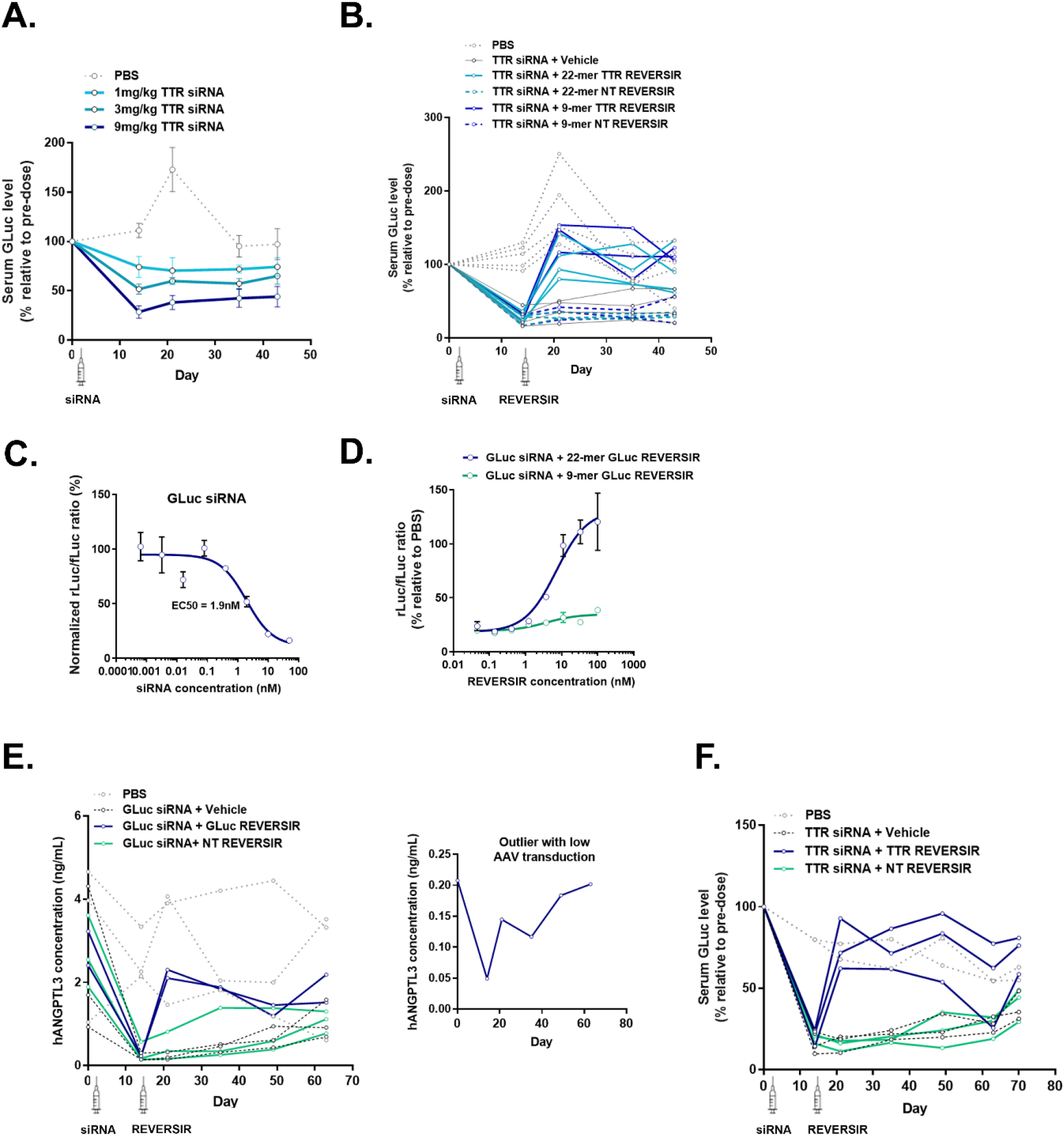
Additional *in vitro* and *in vivo* analyses supporting rAAV regulatory switch leveraging exogenous siRNA and REVERSIR. **(A – C)** Panels represent data from individual animals or additional groups tested as part of the study shown in Fig. 2B – D. rAAV injections, timing of test article dosing, and blood collections were conducted as described in 2B. PBS and 9mg/kg siRNA conditions are identical to those shown in main figure. **(A)** Sustained dose-dependent knockdown of serum GLuc levels in rAAV-injected mice treated with 1,3, and 9mg/kg TTR siRNA as compared to PBS control. **(B)** Averaged data in Fig. 2C presented as spaghetti graph plotting serum GLuc changes over time relative to pre-dose for each individual animal treated with 9mg/kg TTR siRNA followed by 3mg/kg of the specified REVERSIR molecules. **(C)** On-target silencing activity of GLuc siRNA in dual luciferase reporter system. **(D)** Normalized luciferase activity 48h following co-transfection of Cos7 cells with 10nM GLuc siRNA and increasing doses of 22-mer or 9-mer GLuc REVERSIR. **(E and F)** Spaghetti plots showing responses of individual animals that were averaged by condition to generate graphs shown in Fig. 2F and 2G, respectively. **(E - right)** hANGPTL3 concentration over time in one animal that was identified as a significant outlier by Grubb’s test due to low level of rAAV transduction and thus omitted from main figure. All error bars represent s.e.m.

**SUPPLEMENTARY FIGURE 3:**
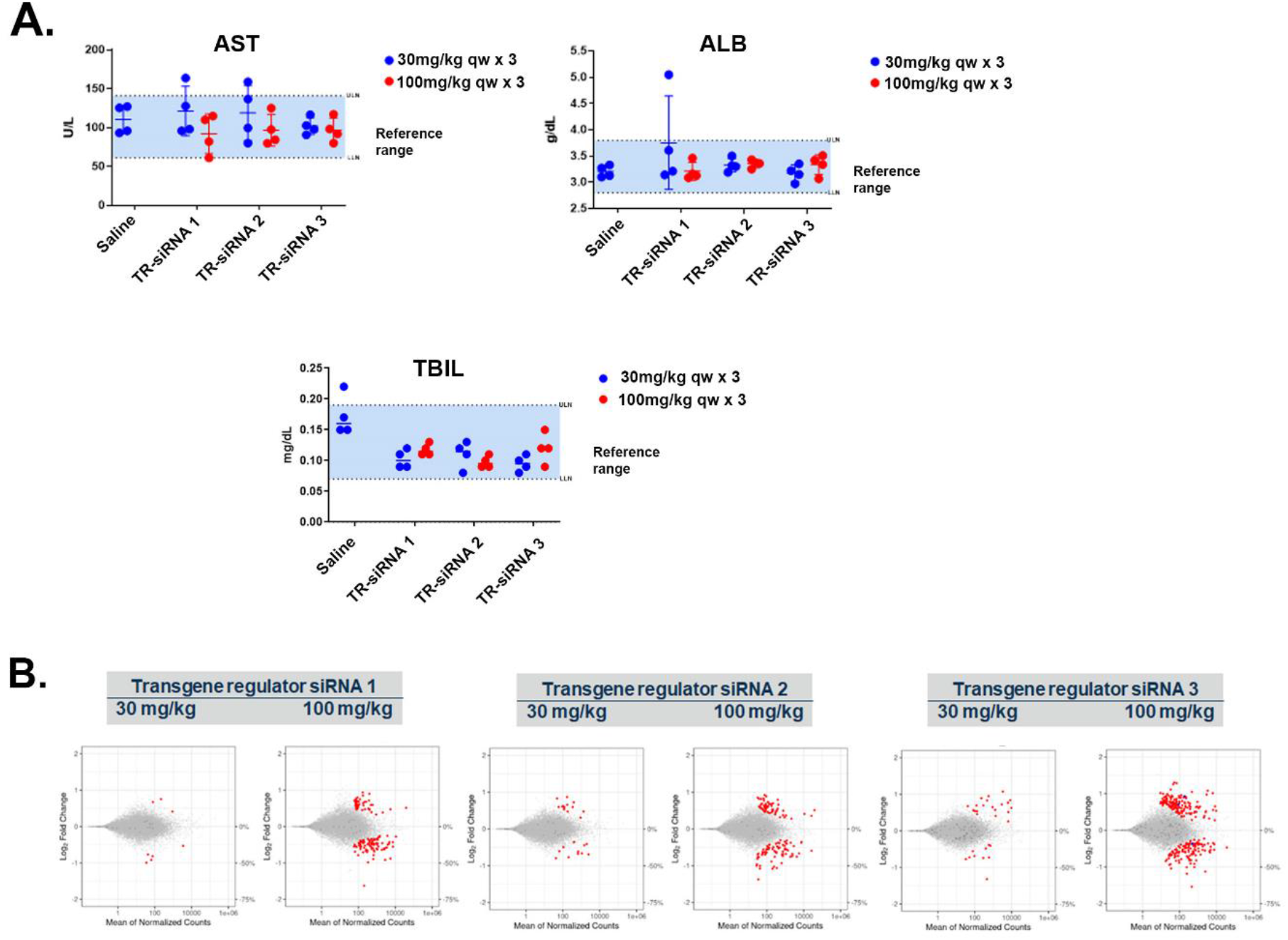
Minimal liver function test (LFT) elevations and transcriptional dysregulation in rat toxicity studies of transgene regulator siRNAs. **(A)** Serum levels of aspartate aminotransferase (AST), albumin (ALB), and total bilirubin (TBIL) at necropsy (D16). Error bars represent s.e.m. **(B**) RNA-seq assessment of rat livers collected 24 h after last dose. Log2 fold change plots (MA plot) represent the average signal from each cohort (*n* = 4). Dots represent individual rat transcripts, their average number of counts and relative change in expression compared to control group dosed with 0.9% NaCl. Gray dots represent gene transcripts that were not found to be differentially expressed following siRNA treatment compared to control. Blue and red dots represent differentially expressed transcripts (false discovery rate < 0.05) with or without a canonical match (8mer, 7mer-m8, 7mer-A1) to the antisense seed region, respectively. Dots in dark gray represent transcripts harboring a canonical seed-matched site but that were not found to be differentially expressed.

**Supplementary Table 1:**
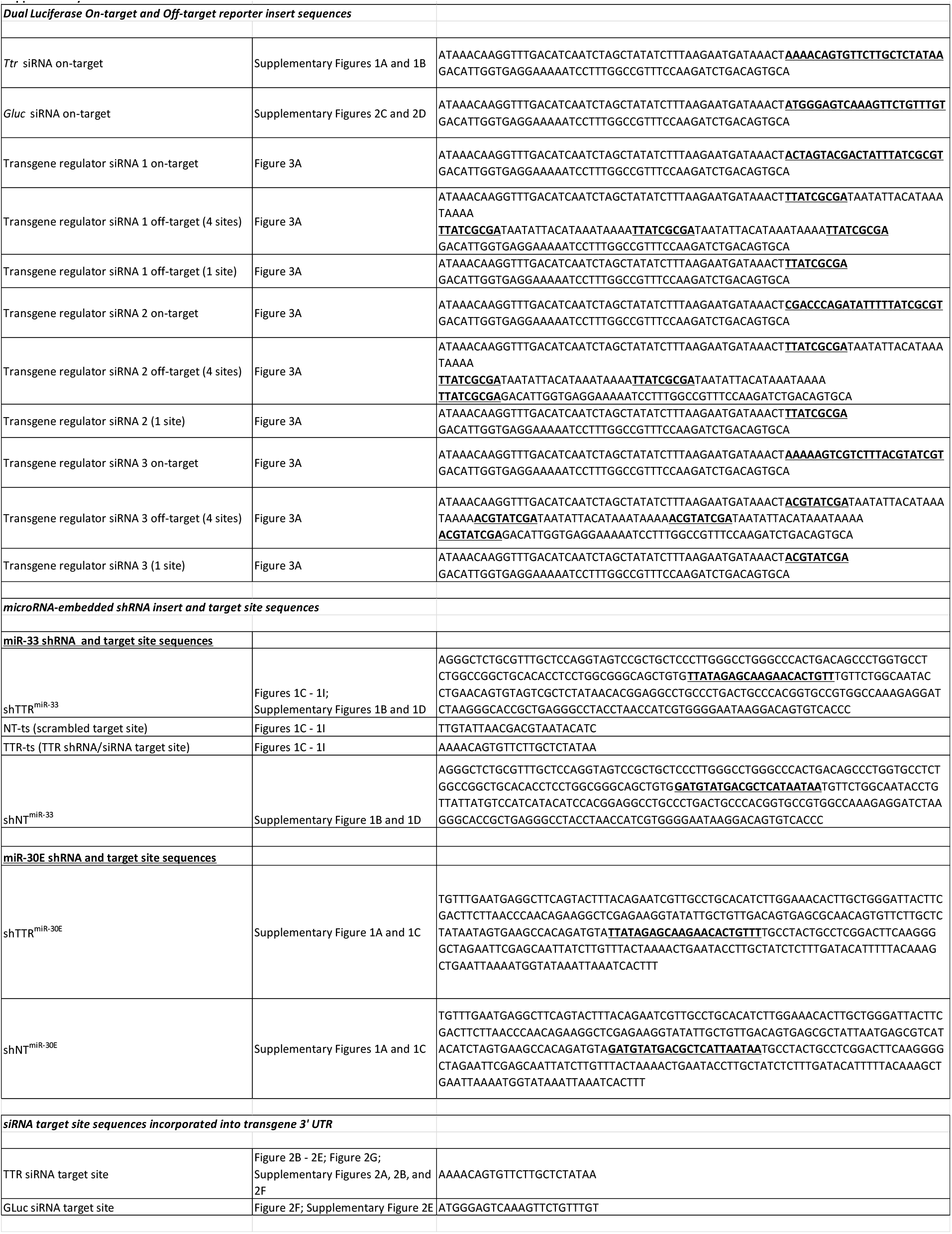
Dual luciferase reporters, shRNA, and RNAi target site sequences used in these studies.

**Supplementary Table 2:** Differential gene expression analysis of *in vitro* RNAseq data from transfection of transgene regulator siRNAs in human and mouse hepatic cells.

| Primary mouse hepatocytes | Number of differentially-expressed genes |  | Strongest DEG<br>(% KD) | Number of<br>2-fold off-targets |
| --- | --- | --- | --- | --- |
|  | Up-regulated | Down-regulated |  |  |
| Transgene regulator siRNA 1 | 13 | 9 | 178% | 0 |
| Transgene regulator siRNA 2 | 0 | 1 | -41% | 0 |
| Transgene regulator siRNA 3 | 44 | 66 | -66% | 5 |
| Hep3B | Number of differentially-expressed genes |  | Strongest DEG<br>(% KD) | Number of<br>2-fold off-targets |
|  | Up-regulated | Down-regulated |  |  |
| Transgene regulator siRNA 1 | 0 | 0 | NA | 0 |
| Transgene regulator siRNA 2 | 0 | 1 | -30% | 0 |
| Transgene regulator siRNA 3 | 0 | 0 | NA | 0 |

